# Effects of zooplankton carcasses degradation on freshwater bacterial community composition and implications for carbon cycling

**DOI:** 10.1101/423772

**Authors:** Olesya V. Kolmakova, Michail I. Gladyshev, Jérémy André Fonvielle, Lars Ganzert, Thomas Hornick, Hans-Peter Grossart

## Abstract

Non-predatory mortality of zooplankton provides an abundant, yet, little studied source of high quality labile organic matter (LOM) in aquatic ecosystems. Using laboratory microcosms, we followed the decomposition of organic carbon of fresh ^13^C-labelled *Daphnia* carcasses by natural bacterioplankton. The experimental setup comprised blank microcosms i.e. artificial lake water without any organic matter additions (**B**), and microcosms either amended with natural humic matter (**H**), fresh *Daphnia* carcasses (**D**) or both, i.e. humic matter and *Daphnia* carcasses (**HD**). Most of the carcass carbon was consumed and respired by the bacterial community within 15 days of incubation. A shift in the bacterial community composition shaped by labile carcass carbon and by humic matter was observed. Nevertheless, we did not observe a quantitative change in humic matter degradation by heterotrophic bacteria in the presence of LOM derived from carcasses. However, carcasses were the main factor driving the bacterial community composition suggesting that the presence of large quantities of dead zooplankton might affect the carbon cycling in aquatic ecosystems. Our results imply that organic matter derived from zooplankton carcasses is efficiently remineralized by a highly specific bacterial community, but doesn’t interfere with the bacterial turnover of more refractory humic matter.

## Introduction

The global carbon cycle is one of the most important biogeochemical processes regulating the climate on our planet (Ward *et al*., 2013). In particular, carbon fluxes between aquatic and terrestrial ecosystems constitute a key component of global biogeochemical cycles (Pace *et al*., 2004; Battin *et al*., 2009; Ward *et al*., 2013). Nowadays, it is well known that a significant part of terrigenous organic matter drains from soils into aquatic ecosystems, especially in the boreal zone (Vachon *et al*., 2017). Freshwaters are considered hotspots of organic matter degradation, sustaining a shorter half-life of organic carbon compared to terrestrial and marine ecosystems (Catalán *et al*., 2015). In freshwaters, organic matter comprises a heterogeneous mixture of different carbon sources with varying degradability. Depending on their degradability by aquatic microbes, the drained terrigenous organic matter is buried to a variable extent in sediments of aquatic ecosystems (Tranvik *et al*., 2009). However, most of the terrigenous (allochthonous) organic carbon is transported into aquatic ecosystems in the form of refractory organic matter (ROM) resulting in a generally higher retention time due to its slow decomposition by aquatic microorganisms (Bianchi, 2011). A major part of this ROM in freshwater ecosystems is represented by humic matter (Rocker *et al*., 2012a).

Bacterial species differ in their response to various sources of carbon resulting in profound implications for aquatic carbon cycling. It has been demonstrated that the availability of organic matter promotes growth of both generalist species, which are able to degrade a wide range of substrates, as well as highly specialized populations degrading specific substrate fractions (Hutalle-Schmelzer *et al*., 2010). Dead zooplankton, which used to be generally neglected in aquatic ecology due to methodological limitations (Tang *et al*., 2009, 2014), is an overlooked and highly abundant source of labile carbon in most freshwater ecosystems. Zooplankton carcasses represent a high quality organic substrate for heterotrophic bacteria due to their relatively low C:N:P ratio as compared to phytoplankton and detritus (Tang *et al*., 2014). Consequently, zooplankton carcasses are “hot spots” of activity of pelagic microorganisms consuming labile organic matter (LOM) as well as ROM (Tang *et al*., 2006; Grossart *et al*., 2007; Elliott *et al*., 2010; Kirillin *et al*., 2012). However, zooplankton carcasses provide not only a carbon source for microorganisms, but also surfaces for attachment. Microorganisms attached to particles are situated in close spatial proximity and can benefit from extracellular degradation enzymes released in the environment (Catalán *et al*., 2015). Thus, attached microorganisms have a higher capacity to degrade polymeric organic matter than their free living counterparts (Grossart, 2010). Consequently, zooplankton carcasses are selecting for specific, but yet uncharacterized microbial communities (Tang *et al*., 2010). The complex LOM of zooplankton carcasses constitutes a valuable source of nutrients and energy for microorganisms, thus implying effects on aquatic carbon cycling, in particular of the more refractory carbon pools. For instance, carcass LOM may induce a “priming effect” and facilitate the degradation of ROM (Bianchi, 2011).

Thus, our primary objective was to investigate consequences on bacterial community composition and carbon cycling in aquatic ecosystems after input of zooplankton carcasses. Since the quality of available organic matter can be a selective force for bacterioplankton community composition (Gómez-Consarnau *et al*., 2012), we tested the hypothesis that nutrient-rich LOM provided by *Daphnia* carcasses selects for generalist bacteria in contrast to C-rich ROM selecting to a larger extent for specialists. In a microcosms experiment, we observed the degradation of ^13^C-labeled carcasses by heterotrophic bacteria from a dystrophic humic bog lake in the presence of indigenous humic matter (treatment *HD*) to track the fate of carcass carbon (Fig. S1). In parallel, we followed three control treatments with either daphnia carcasses (**D**) or humic matter (**H**) as a sole carbon source, and a blank treatment (**B**) containing solely a natural bacterial community. We used optical properties (specific UV absorbance at 254 nm – SUVA_254_, humification index, etc.) and size exclusion chromatography to analyze the influence of carcasses on the dissolved organic matter (DOM) pool and combined it with 16S rRNA gene Illumina amplicon sequencing to characterize the bacterial community composition in detail.

## Results

### Microbial dynamics and community composition

*Daphnia* carcasses showed a rapid decomposition during the first week of incubation, with visible changes in the state of carcasses over time (Fig. S2a-d). At the end of the experiment, the carcasses were still visible as disintegrated parts of the carapace. Dense bacterial colonization of the carcasses was observed (Fig. S2e), while protist grazers or autotrophic organisms were not detected indicating that protozoan grazers and large phytoplankton have been successfully removed by the pre-filtration step. No differences in bacterial counts were observed in the **B** microcosms between the start and the end of the experiment (Table 1). However, a clear increase in bacterial cells counts was observed in *H*, **D** and **HD** microcosms (43%, 654% and 617%, respectively; Table 1). This indicated a higher bacterial growth in the presence of daphnia carcasses and a slower growth on humic matter alone. At the same time, bacterial cell counts did not significantly differ between **HD** and **D** microcosms (paired t-test, p > 0.05, Table 1).

**Table 1.**
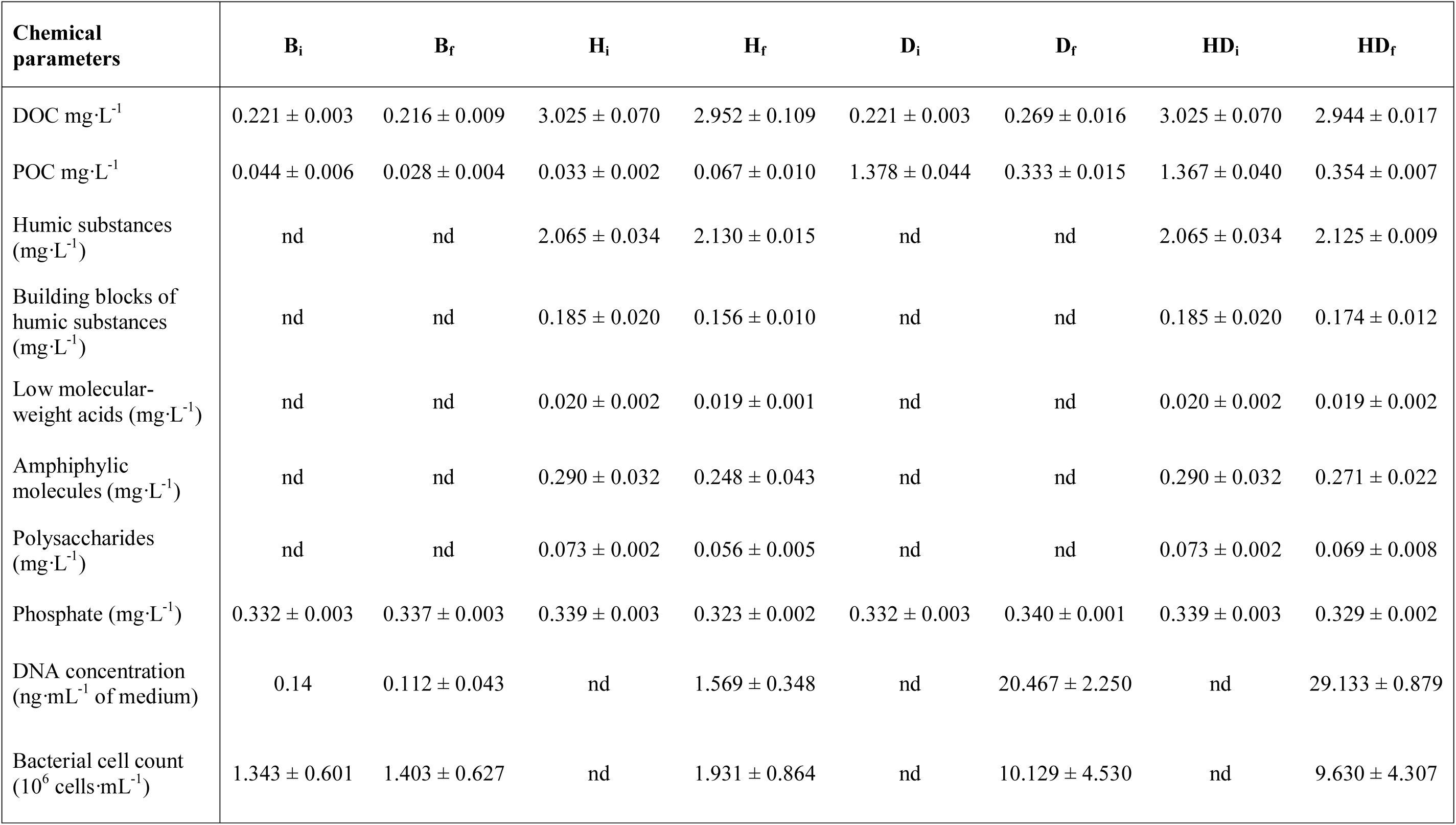
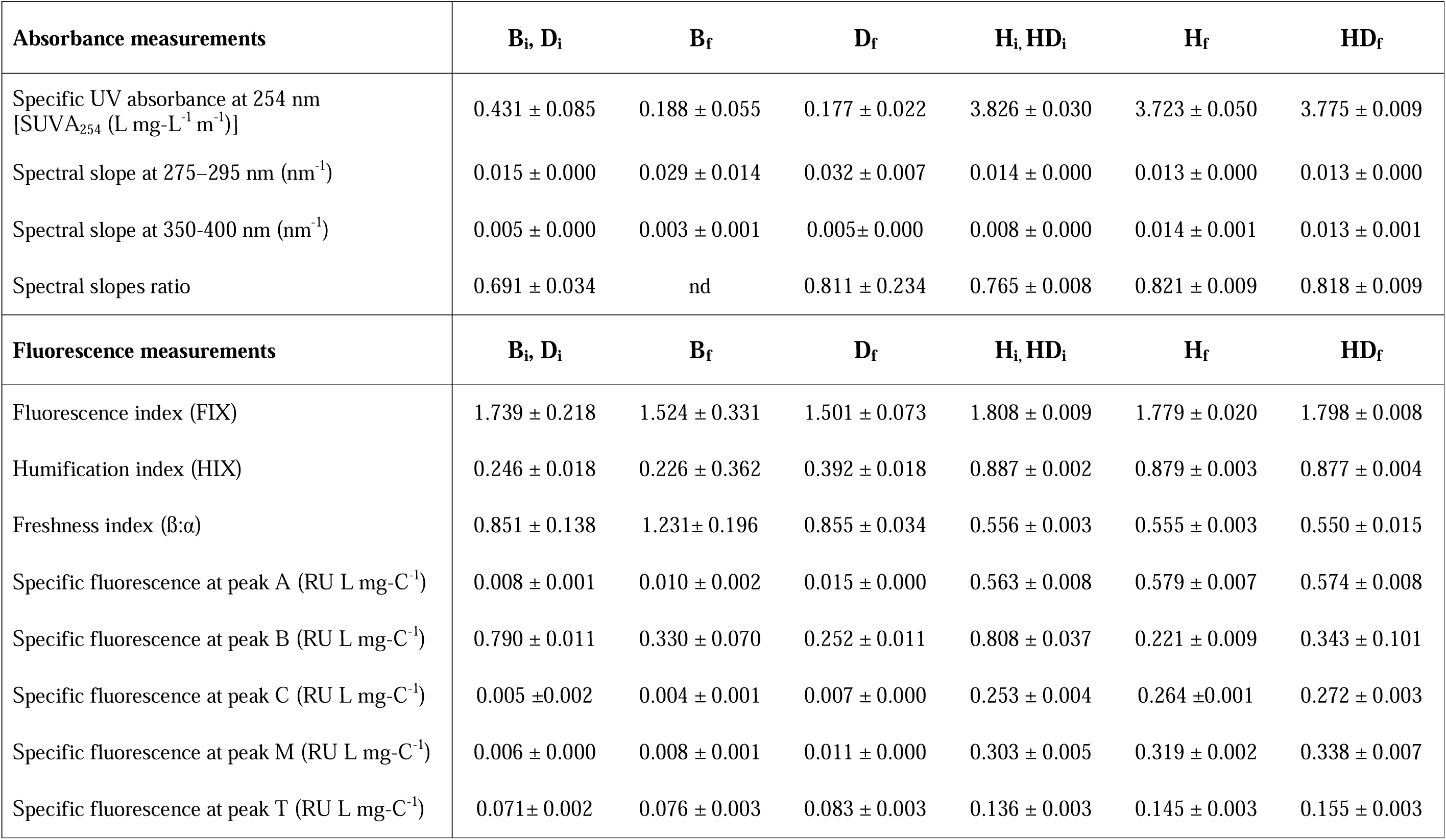
Chemical and optical parameters of experimental microcosms varying in carbon sources: **B_i_** – blank initial, **B_f_** – blank final, **H_i_** – humic matter initial, **H_f_** – humic matter final, **D_i_** – carcasses initial, **D_f_** – carcasses final, **HD_i_** – humic matter + carcasses initial, **HD_f_** – humic matter + carcasses final. Parameters in **D_i_** and **HD_i_** were not measured directly but calculated from parameters in **B_i_** and **H_i_**. Values are given as means of five replicates ± standard errors, except for the initial DNA concentration values obtained from a single measurement. nd = not determined.

After sequencing and performing a quality check for all samples, 595036 reads of 16S rRNA gene fragments were obtained that clustered into 1161 bacterial operational taxonomic units (OTUs). The identified OTUs belonged to 26 known phyla (Fig. 1). In the initial bacterial inoculum from Lake Grosse Fuchskuhle, *Proteobacteria* was the dominant phylum (48% of all sequences), with a high proportion of the class *Betaproteobacteria* (30% of all sequences). By the end of the experiment, the relative abundance of *Proteobacteria* increased in all treatments, especially in carcasses amended microcosms (*HD* and *D*; Fig. 1), with a dominance of the class *Gammaproteobacteria* (62 % and 50 % in **HD** and **D** respectively).

**Fig. 1.**
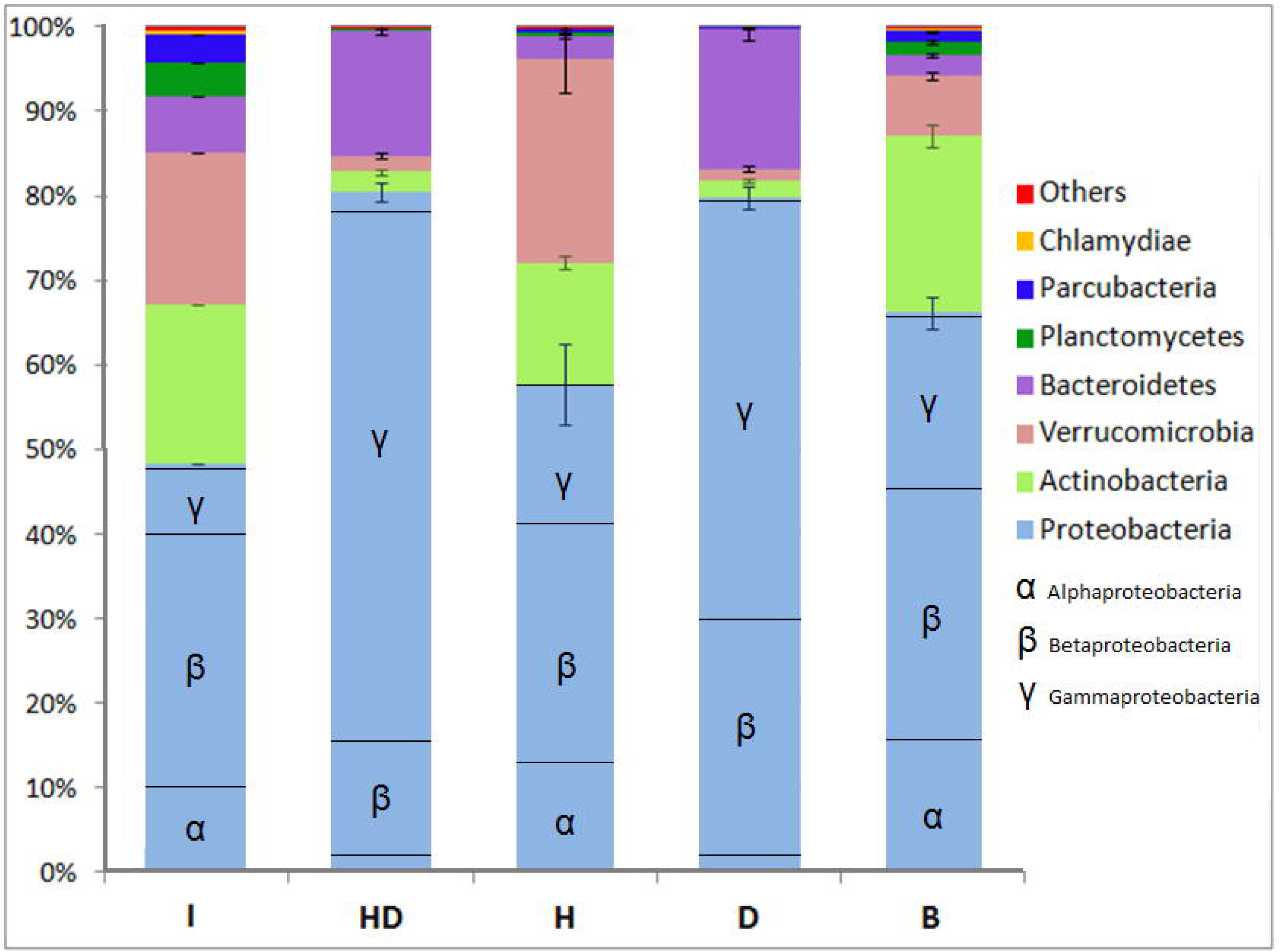
Relative abundance of major bacterial phyla and classes of Proteobacteria in the initial sample (**I**) and the end points of microcosms with additions of humic matter (**H**), *Daphnia* carcasses (**D**), humic matter and carcasses together (**HD**), and blank with no carbon sources (**B**).

At the end of the experiment, the microcosms **HD** and **D** had a lower OTU richness and evenness compared to microcosms without added carcasses (*H* and *B*; Table 2). Microcosms **H** had a lower species richness than **B** microcosms, but showed a higher evenness (Table 2).

**Table 2.**
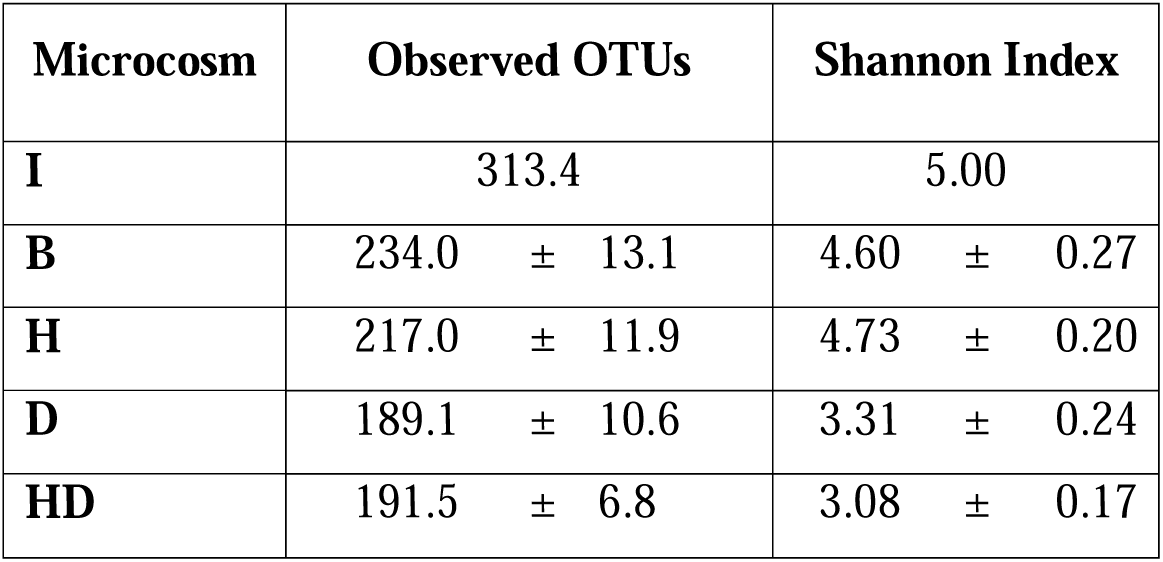
Alpha-diversity of bacterial communities in the microcosms with additions of humic matter (**H**), *Daphnia* carcasses (**D**), humic matter and carcasses together (**HD**), and blank with no carbon sources (**B**). Values are given as means of five replicates ± standard errors, except for the initial inoculate (**I**) obtained from a single measurement, and **B** microcosms (four replicates due to low DNA content in one sample).

In an unconstrained ordination (Fig. 2), all treatments were distinguishable from one another and from the initial inoculum (ANOSIM, R = 0.835, p = 0.001). The OTUs accounting for most of the difference between the treatments were identified using a SIMPER test (Table 3). The relative abundance of the most influential OTU (OTU1, *Pseudomonas* sp.) was significantly different between the start and the end of the incubation for all treatments (ANOVA F_4,15_ = 102.89, p < 0.001). The distribution of different OTUs in all treatments is discussed in more details in the Supplementary Information.

**Fig. 2.**
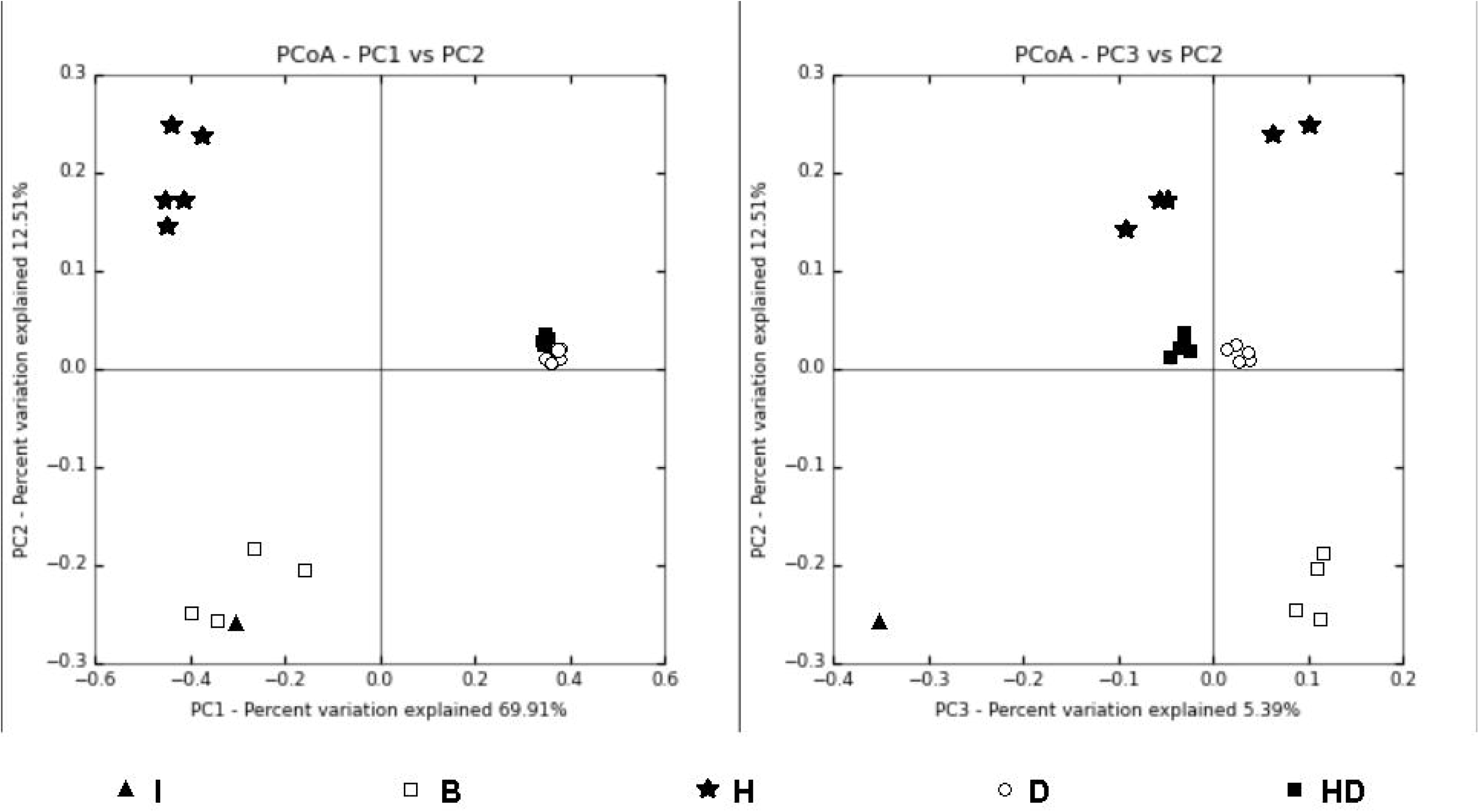
Principal coordinate analysis of the initial sample (I) and the end points of microcosms with additions of humic matter (**H**), *Daphnia* carcasses (**D**), humic matter and carcasses together (**HD**), and blank with no carbon sources (**B**), based on Bray-Curtis community similarity, calculated as relative abundance of operational taxonomic units (OTUs).

**Table 3.**
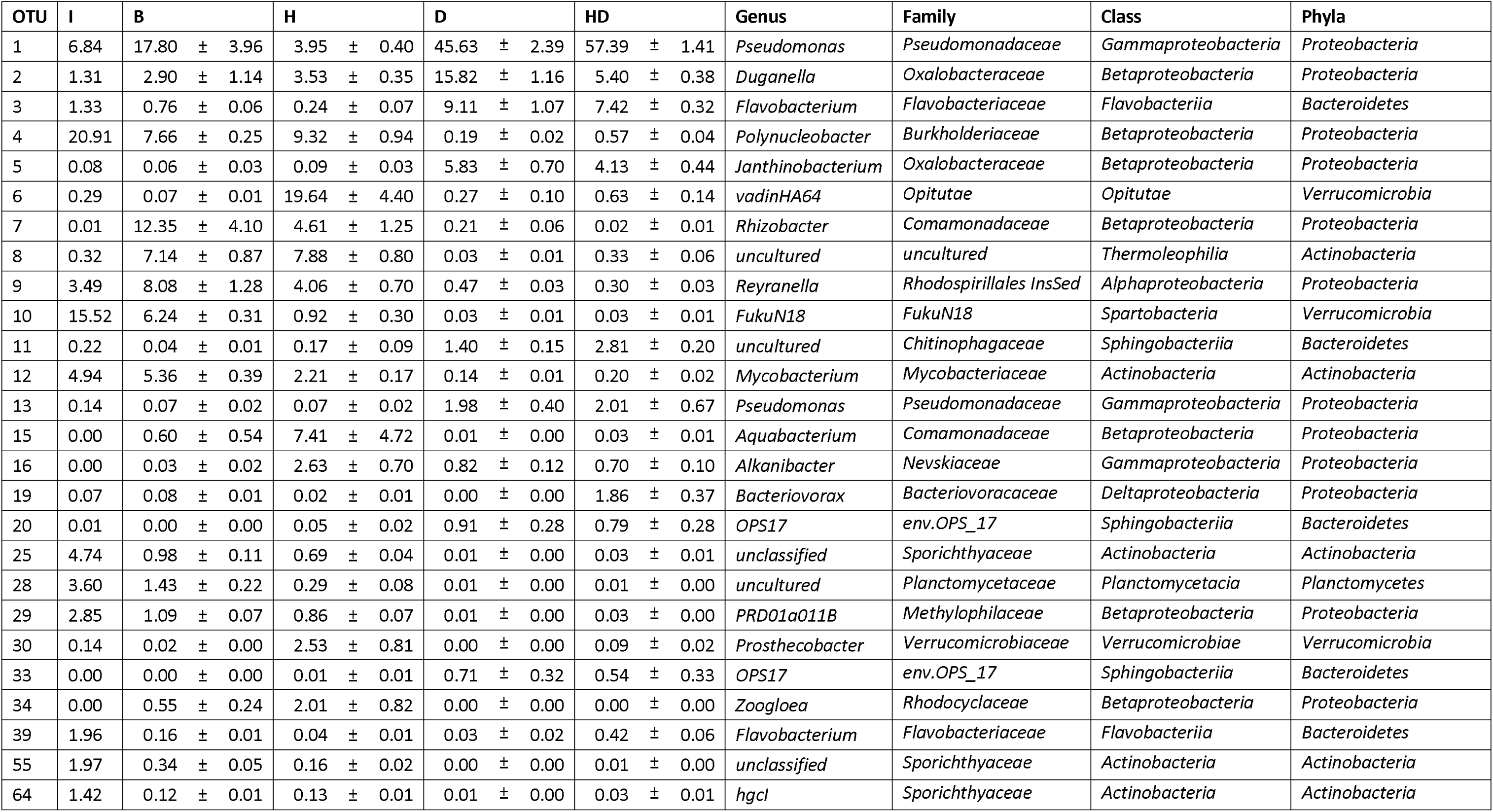
Abundances (in %) of the quantitatively prominent OTUs contributing the most to the dissimilarity among the inoculum sample (**I**) and the end points of the microcosms with additions of humic matter (**H**), *Daphnia* carcasses (**D**), humic matter and carcasses together (**HD**), and blank with no carbon sources (**B**). Values are given as means of five replicates ± standard errors, except for the inoculum sample obtained from a single measurement, and **B** microcosms (four replicates due to low DNA content of two samples which were pooled).

### DOM composition and fate of carcass carbon

We aimed to test for the “priming effect” by comparing the predicted total organic carbon degradation rate (ΔTOC) in **HD** microcosms, calculated from ΔTOC in the control microcosms *D*, *H*, and *B*, to the measured ΔTOC in **HD** microcosms (see Supplementary Methods for more details). The predicted ΔTOC was -1.015 ± 0.115 mg L^−1^ and did not differ significantly (paired t-test, p > 0.05) from the ΔTOC measured in **HD** microcosms (-1.094 ± 0.057 mg L^−1^).

During carcass decomposition (in **HD** and **D** microcosms only), the particulate organic carbon (POC) decreased approximately four-fold compared to the initial values (Table 1). Nevertheless, no significant differences in concentrations of dissolved organic carbon (DOC), high- and low-molecular weight substances (HMWS and LMWS, respectively) or humic substances were found throughout the experiment in any treatment (paired t-tests, p > 0.05 in all tests, Table 1). However, in ROM supplemented microcosms (*HD* and *H*) we observed trends of decreasing concentrations in polysaccharides, amphiphilic molecules, and building blocks of humic substances (Table 1).

The SUVA_254_, spectral slopes and optical indices values were not different between the humic-amended microcosms **H** and **HD** (Table 1), and they did not differ between **D** and **B** microcosms (Table 1). An exception was the freshness index, being an indicator for recently produced DOM (Hansen *et al*., 2016), which was lower in **D** compared to **B** microcosms. Expectedly, the ratio of peakA to peakT, which is known as an indicator of the ratio of humic-like (recalcitrant) to freshly produced (labile) organic matter (Hansen *et al*., 2016), was higher in **H** microcosms compared to **HD** (Wilcoxon test, p-value = 0.03).

The carcasses had an average ^13^C/^12^C ratio of 0.168 ± 0.004 (δ^13^C = 13945.3 ± 303.9‰). According to this specific signature, the amount of processed carbon originating from the carcasses was computed in all carcass-containing microcosms (i.e. **HD** and **D**). In **HD** microcosms, 72.8% of POC and 2.2% of DOC originated from daphnia carcasses, against 82.1% of POC and 21.1% of DOC for the **D** microcosms (Fig. 3).

**Fig. 3.**
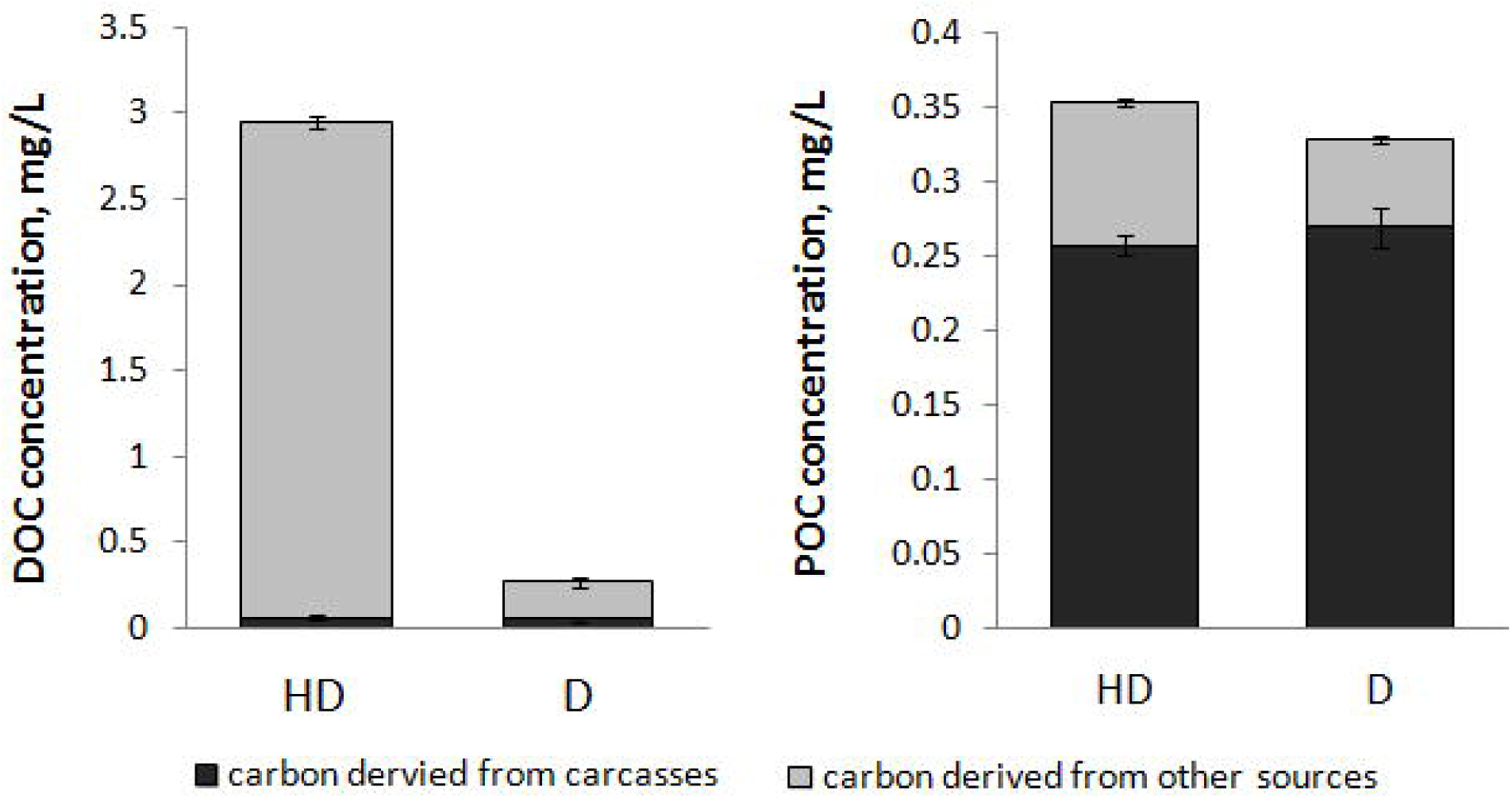
Carbon fractions originated from daphnia carcasses at the end of the experiment in pools of dissolved (DOC) and particulate organic carbon (POC) as well as in dissolved CO_2_ in microcosms with humic matter and *Daphnia* carcasses (**HD**) and carcasses only (**D**). The area size of diagrams is relative to carbon concentration.

The bacterial respiration, measured as the increase in CO_2_ and normalized to the background respiration (i.e. respiration from the blank microcosms **B**) was higher in **D** microcosms compared to **H** but lower compared to **HD** microcosms (Fig. S3). The respiration per amount of initially added carbon and normalized to the background respiration was higher in **HD** microcosms compared to **H** but lower compared to **D** microcosms, and was used to confirm the labile character of the organic matter originating from the zooplankton carcasses. All differences in CO_2_ concentrations between the microcosms were significant (1-way ANOVA F = 241.6, p < 0.001; Tukey post hoc test p < 0.01 for all pairs).

To test for the priming effect, the predicted ^13^C/^12^C ratio in the respired CO_2_ of **HD** microcosms was calculated from the values in the control microcosms *D*, *H*, and *B*, and compared to the measured ^13^CO_2_/^12^CO_2_ ratio in **HD** microcosms (see Supplementary Methods for more details). The predicted ^13^CO_2_/^12^CO_2_ ratio for **HD** (0.078 ± 0.003) did not differ significantly (paired t-test, p > 0.05) from the measured value (0.073 ± 0.003). In **HD** microcosms, 87.8 ± 6.3 % of respired CO_2_ originated from zooplankton carcasses when normalized to the background respiration of microcosms **B**.

### Interactions between microbial community and DOC quality

The interactions between bacterial community composition and DOC quality in microcosms **HD**, *H*, and **D** revealed specific patterns in bacterial substrate preferences (Fig. 4). The connections between bacterial genera and DOC qualities significantly differed between microcosms **H** and **D** (Fig. 4a). Thus, bacteria positively interacting with DOC concentration as well as humification and fluorescence indices are the ones thriving in the presence of humic matter (Fig. 4a). On the contrary, bacteria negatively associated with these parameters are favored by carcasses (Fig. 4a). Similarly, genera positively correlated with DOC concentration, fluorescence index and A/T peak ratio (Fig. 4b, comparing microcosms **HD** and **D**) seem to be favored by humic matter when daphnia carcasses are available. However, those genera negatively correlated with these parameters are suppressed by humic matter in the presence of carcasses.

**Fig. 4.**
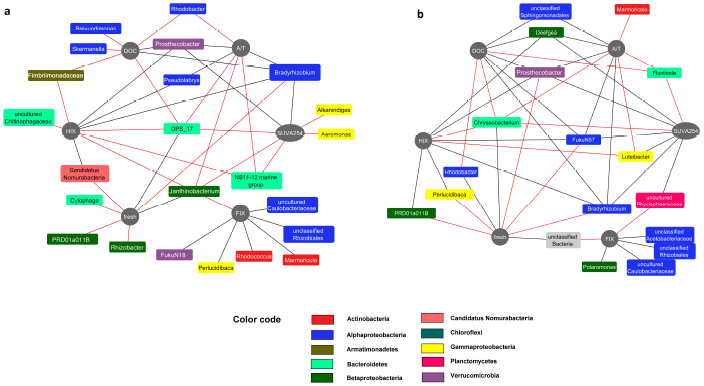
Network of bacterial genera connected with major DOM parameters, significantly different in pairs of microcosms: a – microcosms **H** – humic matter, and **D** – carcasses; b – microcosms **HD** – humic matter + carcasses, and **D** – carcasses. Nodes that interact positively are connected by solid black edges, nodes connected by dashed edges have negative interactions. Legend: DOC – concentration of DOC; HIX – humification index, FIX – fluorescence index, fresh – freshness index, SUVA254 – Specific UV absorbance at 254 nm, A/T – ratio between the specific fluorescence at peak A (excitation 240-260 nm/emission 400-500 nm, UVC humic-like fluorescent component) and peak T (excitation 270-285 nm/emission 340-380 nm, tryptophan-like fluorescent component).

## Discussion

The main objective of this study was to track the degradation of zooplankton carcasses, as a so far largely neglected but common and labile carbon source (Tang *et al*., 2014). Our study reveals selection of defined bacterial communities in the presence of carcasses strongly related to the specific DOM quality released from carcasses. However, carcass-induced availability of LOM and related shifts in bacterial community composition did not result in significant changes in the turnover of the added ROM pool. Consequently, our data do not support a “priming effect” (Bianchi *et al*., 2011) of refractory humic matter removal in the presence of relatively labile carbon from *Daphnia* carcasses. Nevertheless, the bacterial community composition was greatly affected by the presence of carcass carbon. Thus, our study adds new quantitative and qualitative data on bacterial carbon utilization related to changes in the community composition induced by changes in substrate quality, i.e. addition of zooplankton carcasses, and adds new insights in microbial-organic matter interactions.

### Bacterial community composition depending on carbon source

As outlined above, we did not measure any quantitative changes in organic matter degradation between **HD** microcosms and the predicted values calculated based on the parameters of single-carbon source microcosms **H** and **D**. Consequently, the presence of LOM from daphnia carcasses did not change the degradation of humic matter but rather influenced the bacterial community.

According to our data on beta-diversity of bacterial assemblages (Fig. 2), the type of treatment strongly affected the bacterial community composition in each microcosm. In microcosms with no extra organic matter addition (**B**), the bacterial community remained similar to the initial inoculum indicating that experimental changes in environmental conditions did not modify the bacterial community composition drastically (Fig. 2). Overall, the addition of carcass LOM was the main driver of the bacterial community composition in the **D** and **HD** microcosms (Fig. 2). Bacterial taxa introduced into the microcosms with the carcasses may also have an effect on bacterial community composition and richness. However, according to literature data, zooplankton carcasses are not primarily decomposed by their native-associated bacterial communities, but rather by ambient bacteria (Bickel and Tang, 2010). This notion is also reflected by the fact that bacteria richness is the lowest in the **HD** and **D** treatments. Thus, in natural waters with a high amount of dead zooplankton, carcasses can be a primary factor driving bacterioplankton community composition with potential effects for carbon cycling. In many lakes with an “inverted” biomass pyramid, zooplankton biomass is higher than phytoplankton biomass (Heathcote *et al*., 2016). We suppose that the same pattern for dead biomass would indicate that zooplankton and not algal LOM, which is usually in the researcher’s focus (Hoikkala *et al*., 2016; Landa *et al*., 2016), could be the major driver for bacterioplankton community composition in such ecosystems. In lakes with a “normal” biomass pyramid, zooplankton may still play an important role for determining bacterial community composition in the occasional events of mass zooplankton mortality (Tang *et al*., 2014). It would be interesting to test this presupposition in further studies. The co-presence of ROM also contributed to specific bacterial communities by selecting for a number of specific OTUs. The observed, significant difference in bacterial community composition between all treatments points to a pronounced effect of substrate quality on bacterial community composition and might result in functional differences.

The prevalence of *Betaproteobacteria* in the initial inoculum and in microcosms **B** and **H** (Fig. 1) was in accordance with previous studies on Lake Grosse Fuchskuhle (Grossart *et al*., 2008; Hutalle-Schmelzer *et al*., 2010). Indeed, *Betaproteobacteria* are among the most numerous bacteria in the upper layers of freshwater lakes, in particular of peat bog lakes (Newton *et al*., 2011). Moreover, the dominance of *Gammaproteobacteria* in carcass-amended microcosms (Fig. 1) was also expected according to previous studies (Tang *et al*., 2009; Shoemaker and Moisander, 2015). *Gammaproteobacteria* include many species with a copiotrophic lifestyle that can grow faster than the average lake bacterioplankton, especially under nutrient-rich conditions as can be found on carcasses (Newton *et al*., 2011). Interestingly, a more distinct community pattern emerged in the different microcosms when taking the level of individual OTUs into account. This indicates a close relationship between organic carbon quality and bacterial community structure (Attermeyer *et al*., 2014, 2015).

### Links between organic matter quality and microbial community composition

A number of uncultivated bacterial taxa belonging to the order *Sphingobacteriales* were positively selected solely in the presence of carcasses only (Fig. 4a). Many members of *Sphingobacteriales* express chitinolytic activity (Kämpfer, 2015), but no information is available in particular about the ecological role of the uncultivated representatives found in the present study. High abundances of the NS11-12 marine group were previously associated with an increase in chlorophyll *a* concentration (Meziti *et al*., 2015) and number of particles, while it is negatively correlated with nitrate concentration (Henson *et al*., 2016). Moreover, the uncultured bacterial group OPS 17 was previously found not to respond to terrestrial DOM additions (Lindh *et al*., 2015).

Another group of bacteria favored by zooplankton carcass LOM were ubiquitous chemoorganotrophs belonging to genera *Brevundimonas* and *Aeromonas* (Segers *et al*., 1994).

Among the bacteria favored by humic matter (Fig. 4b), genera involved in nitrogen fixation (*Bradyrhizobium*, *Rhizobacter*, unclassified *Rhizobiales*) (Kuykendall, 2005; Goto, 2015), organic pollutants degraders *Rhodococcus* (Bell *et al*., 1998), unclassified *Sphingomonadales* and methylotrophs (uncultured strain PRD01a011B from *Methylophilaceae* (Doronina *et al*., 2014)) were detected. To a large extent the same bacterial genera were favored by humic matter regardless whether zooplankton carcasses were present. In contrast, bacteria suppressed by humic matter in the presence of carcasses were almost exclusively chemoorganotrophic generalists (Johansen *et al*., 2005; O’Sullivan *et al*., 2005; Song *et al*., 2008; McBride, 2014; Evtushenko, 2015).

Besides selecting for certain bacterial populations, zooplankton carcasses strongly decreased the species richness and evenness of the bacterial community (Table 2). Therefore it appears that the availability of a high quality and abundant LOM source can reduce the biodiversity by favoring a small number of copiotrophs dominating the community: in the present study about half of all sequences in the **HD** and **D** microcosms belonged to a single OTU (OTU 1, *Pseudomonas sp*.). In a similar study of Blanchet et al. (2017), the bacterial diversity was not affected by amino acid additions, possibly because free amino acids are simple compounds which can be consumed simultaneously by many members of the community (Trusova et al., 2012). Thus, it is solely by providing a surface for attachment in combination with a complex but yet labile carbon source that zooplankton carcasses are changing bacterial communities towards a less diverse, more uneven and more copiotrophic community.

### Microbial carcass decomposition in relation to carbon quality

An abrupt increase of LOM availability following mass zooplankton mortality commonly observed in natural waters (reviewed by Tang et al. 2014), leads to a substantial input of both DOC and POC. In agreement with a previous study (Tang *et al*., 2006), *Daphnia* carcasses lost their mostly labile internal tissues rapidly, whereas the chitin-based carapace was more resistant to dissolution and microbial decomposition (Fig S2).

However, in our study the leached fraction was rapidly consumed by numerous ambient bacteria (Table 1), and did not increase the DOC concentration significantly in the microcosms with zooplankton carcasses (**D** and **HD**). This statement is supported by our observation that adding 1.334 ± 0.038 mg C L^−1^ with *Daphnia* carcasses resulted in only 0.055 ± 0.011 and 0.065 ± 0.010 mg C L^−1^ of carcasses-derived DOC in **D** and **HD** microcosms, respectively (Fig. 3).

On the other hand, the chitin-based structure of the carapace was only partially degraded and also used a surface for attachment. At the end of the experiment, 0.270 ± 0.014 and 0.258 ± 0.007 mg C L^−1^ originating from daphnia carcasses (19-20% of the initial quantity) remained in the POC fraction of the **D** and **HD** microcosms, respectively (Fig. 3), mainly represented by the remaining carapace and the bacterial biomass (Fig. S2). This is further confirmed by the finding that bacterial taxa degrading chitin were greatly favored in the presence of carcasses at the end of the incubation (Fig. 4). The difference between non-carcass-derived POC in microcosms **D** and **HD** (0.059 ± 0.003 and 0.096 ± 0.003 mg L^−1^, respectively; Fig. 3) presumably occur due to humic matter aggregation converting DOC into POC. This assumption is supported by a two-fold POC increase in the **H** microcosms compared to the initial value (Table 1).

In our experiment we used 40 *Daphnia* carcasses per liter, a high but still natural value (Dubovskaya *et al*., 2003). Most of the organic matter originating from carcasses was respired by the bacterial community within the two weeks of incubation. Therefore, in natural systems, the considerable amount of LOM released by zooplankton carcasses (Tang *et al*., 2014), can directly affect the functioning of the ecosystem by accelerating microbial carbon turnover and respiratory carbon losses to the atmosphere at short time scales. The more recalcitrant part of the carapace may persist for a longer time, and eventually escapes the water column to be further processed in the sediments (Tang *et al*., 2014), being also important for carbon sequestration. Consequently, the balance between microbial degradation of zooplankton carcasses and organic matter storage in sediments has a great influence on the aquatic carbon cycle.

### Priming, a concept under debate in aquatic sciences

Humic matter was chosen as a recalcitrant carbon source for its ubiquity in aquatic ecosystems and as an important part of the carbon pool in the global carbon cycle. Humic matter can represent up to 80% of the total DOM in freshwaters (Rocker *et al*., 2012a). Although humic matter is considered as recalcitrant, it can at least partially be decomposed by bacteria (Hutalle-Schmelzer *et al*., 2010; Rocker *et al*., 2012a; Kisand *et al*., 2013). Furthermore, the degradation of humic acids by marine and estuarine bacterial communities seems to be favored by specific environmental conditions, e.g. along a salinity gradient (Rocker *et al*., 2012a; Rocker *et al*., 2012b; Kisand *et al*., 2013). Consequently, recalcitrance and lability of organic matter are not *per se* intrinsic chemical characteristics (Schmidt *et al*., 2011), and may only account for specific environmental settings (Bianchi *et al*., 2015).

In our experiment, we incubated daphnia carcasses and humic matter in different combinations to test for a priming effect of bacterial degradation of ROM induced by the addition of carcass LOM. The addition of a mixture of natural humic matter and *Daphnia* carcasses resulted in the degradation of organic matter and an isotopic ratio of the respired CO_2_ similar to what we predicted based on our linear addition model with humic matter or carcasses as the sole carbon source. Moreover, based on DOM characterization (Table 1), the natural humic matter from Lake Grosse Fuchskuhle was only little degraded by bacteria, irrespective of LOM addition via daphnia carcasses. Consequently, and in agreement with previous studies (Bengtsson *et al*., 2014; Catalán *et al*., 2015; Dorado-García *et al*., 2015), we could not detect any quantitative changes in bacterial ROM degradation when using carcass LOM as a potential primer (Table 1).

Recently, a number of studies have investigated the prevalence of a priming effect in aquatic ecosystems (e.g., van Nugteren *et al*., 2009; Guenet *et al*., 2013; Kuehn *et al*., 2014; Steen *et al*., 2015). Various types of ecosystems (marine, lentic, lotic) and habitats (pelagic, hyporheic, sediments) have been tested, as well as different sources of LOM (carbohydrates, algae leachate, gastropod mucus, etc.) and ROM (terrestrial plant tissues, lignocellulose, humic matter, etc.) have been used. Although some authors have found support for ROM priming by more labile organic matter, mainly of algal origin (van Nugteren *et al*., 2009; Guenet *et al*., 2013; Hotchkiss *et al*., 2014; Bianchi *et al*., 2015; Gontikaki *et al*., 2015), others did not reveal any evidence for a positive priming effect (Bengtsson *et al*., 2014; Catalán *et al*., 2015; Dorado-García *et al*., 2015; Blanchet *et al*., 2017), or even found a negative priming effect (Gontikaki *et al*., 2013) with ROM being decomposed slower in the presence of a labile carbon source. Thus, it appears that the absence or presence of the priming effect may strongly depend on specific environmental or experimental conditions, which may also explain the absence of humic matter degradation in our **HD** treatments using daphnia carcasses LOM as the primer.

### Limitations and applications of the study

The degradation of ROM (such as humic matter) is a combination of two main processes: microbial and photochemical decomposition (Amado *et al*., 2015). The most efficient humic matter microbial degraders in aquatic systems are fungi (Grinhut *et al*., 2007). Generally, fungi have a higher capacity then bacteria to synthesize the extracellular oxidative enzymes involved in ROM degradation and thus more readily and successfully initiate humic matter degradation (Rojas-Jimenez *et al*., 2017). In contrast, bacteria join the process later as degraders of humic matter metabolites (Grossart and Rojas-Jimenez, 2016; Rojas-Jimenez *et al*., 2017). Due to our pre-filtration step to avoid the presence of protozoan grazers and large phytoplankton as has been frequently done in similar incubation experiments (Fonte *et al*., 2013; Guenet *et al*., 2013; Attermeyer *et al*., 2014, 2015; Blanchet *et al*., 2015), fungi, which could potentially constitute an important component of the aquatic priming effect, were removed.

Photochemical degradation can break down/oxidise recalcitrant DOM compounds, such as fulvic and humic substances, into more labile molecules (Spencer *et al*., 2009; Stubbins *et al*., 2010). It is likely that in natural systems photochemical and microbial degradation work synergistically and thus contribute to the priming effect. On the other hand, UV radiation can also affect biodegradability of LOM in the presence of humic matter (Tranvik and Kokalj, 1998). Our study was conducted in the dark as in other priming effect studies (e.g. Bianchi *et al*., 2015; Catalán *et al*., 2015) to avoid photosynthesis which could have obstructed the detection of differences in carbon oxidation between treatments and controls. Thus, it is clear that photodegradation of humic substances was not taken into account in the present study and might have reduced our possibilities to measure a positive priming effect. However, our study is directly applicable to the situation when the zooplankton carcasses are sinking into the pelagic zone at depths with very low light penetration.

The incubation temperature in our experiment was relatively high (20°C) and not typical for deep waters. Nevertheless, if temperature might have a strong influence on bacterial community composition and its activity (Adams *et al*., 2010), a lower temperature only slows down the degradation process but not the biodegradability of carcasses. Therefore, in natural systems, the potential of zooplankton carcasses to release a consequential amount of carbon in the atmosphere and to select for specific bacterial community might happen over longer time scales than the ones observed in our experiment.

Another factor that can affect DOM degradation is the oxygen concentration, which usually decreases sharply with depth in the pelagic zone of humic lakes. In our experimental microcosms we maintained 100% oxygen saturation due to mixing with headspace air. However, Tang *et al*. (2006) observed only a small difference between microbial communities decomposing carcasses in anaerobic vs. aerobic conditions, presumably because zooplankton carcasses represent anoxic microenvironments even when oxygen is abundant in the surrounding waters (Glud *et al*., 2015).

Although the experimental set-up might have its limitations, our findings are relevant to natural environments such as the metalimnion of humic lakes and other deep water layers where the maximum percentage of dead zooplankton is found (Dubovskaya *et al*., 2018).

## Conclusions

We observed a pronounced change in bacterial community composition in microcosms induced by the addition of *Daphnia* carcasses and humic matter. While the quality of both added carbon sources played a role, zooplankton carcasses were the major driver of bacterial community assembly. Even though no priming effect was detected, it is critical to continue studies on ROM degradation in the presence of zooplankton carcasses to better understand microbial dynamics and thus potential changes in organic matter fluxes in freshwater ecosystems. In events of mass zooplankton mortality, the water column can be loaded with a considerable amount of carcass-derived LOM. By decreasing the bacterial diversity and selecting specialized bacteria, zooplankton carcasses potentially have further and so far likely unknown consequences on microbial dynamics and carbon fluxes. Indeed, our results suggest that a significant part of zooplankton carcasses is respired by bacteria. Therefore, our study provides evidence that quantifying the implication of zooplankton carcasses in the functioning of aquatic ecosystems is of primordial importance to understand the amount of carbon produced in and released from freshwaters.

## Experimental Procedures

### Experimental setup

The experiment was conducted in laboratory microcosms, which were set up in 1L acid-washed and muffled (4h, 450 °C) glass bottles half-filled with artificial lake water. The microcosms were inoculated with a concentrated bacterial community from the acidic bog lake Grosse Fuchskuhle (Northeastern Germany, 53°06’N 12°59’E; more detailed information is provided in the Supplementary Methods). Microcosms were sealed with PTFE-coated silicone septa, placed on a roller apparatus (Wheaton, USA), and incubated for 15 days in the dark at 20°C. The duration of the experiment, irradiance, temperature, and the rolling mode were chosen to fall within the range of natural conditions of sinking zooplankton carcasses in a water column (Tang *et al*., 2009, 2014).

One set of microcosms (**HD**) was amended with humic matter and ^13^C-labeled *Daphnia* carcasses (Fig. S1). The detailed description of the amendments including their preparation is available in the Supplementary Methods. Carcasses control microcosms (**D**) were amended solely with ^13^C-labeled *Daphnia* carcasses in the same quantity and from the same batch of *Daphnia* as in **HD** microcosms. Humic matter control microcosms (**H**) were amended with humic matter only in the same quantity as in **HD** microcosms (Fig. S1). Further, blank microcosms (**B**) with no added organic matter were included to determine whether the bacterial community was capable of growing without the extra addition of organic carbon (Fig. S1). Each experimental treatment was conducted in five replicates.

### Bacterial counts and community composition

Microbial abundances were determined after filtration of 5 mL of water through Nucleopore track-etched membrane filters with a pore size of 0.2 μm (Whatman, UK). Then, samples were stained with 4’,6-diamidino-2-phenylindole (DAPI) to monitor cell numbers using an imaging system linked to a Leica epifluorescence microscope. Pictures were taken from 30-50 fields and abundances were determined using the CellC software (Tampere University of Technology, Finland, https://sites.google.com/site/cellcsoftware/).

For DNA extraction, 150 mL of water was filtered through 0.22 μm GVWP filters (Millipore, Germany) and total DNA was extracted according to a modified protocol described by Nercessian *et al*. (2005) (see Supplementary Methods for details). DNA concentrations were determined with a Quantus fluorometer (Promega, USA), following the manufacturer’s instructions. PCR, library preparation and sequencing was done by LGC Genomics (Berlin, Germany). Briefly, the V3-V4 region was amplified using primers 341F-785R (Klindworth *et al*., 2013), followed by library preparation (2x300 bp) and sequencing on a MiSeq Illumina platform. Sequences were quality checked and analyzed using Mothur v1.37.6 (Schloss *et al*., 2009), see Supplementary Methods for the detailed workflow. The sequence data was deposited in Genbank under the following accession number: PRJNA418906.

### Organic carbon concentration and composition

Directly after sampling, water was passed through pre-combusted (4 h, 450°C) GF75 filters (Advantec, nominal pore size of 0.3 μm). Particulate organic carbon (POC) collected on the filters was measured with an Eltra SC 800 (Eltra, Germany). One subsample of filtrate was processed directly to measure the DOC concentration with a TOC-VCPH (Shimadzu, Kyoto, Japan) as well as the phosphate concentration using a FIAstar 5000 (Foss, USA). Total organic carbon (TOC) concentration was estimated by summing up the DOC and POC concentrations.

Another subsample of filtrate was stored at 4°C for three weeks prior analysis with liquid chromatography - organic carbon detection – organic nitrogen detection (LC-OCD-OND, DOC Labor, Germany). The LC-OCD-OND allows distinguishing between high- and low-molecular weight substances (HMWS and LMWS, respectively) and humic substances (Huber *et al*., 2011). Nevertheless, its sensitivity is not sufficient for analyzing samples with low DOC concentration (<0.2 mg L^−1^). That was the case for the **D** and **B** microcosms which were therefore not analyzed by LC-OCD-OND.

### Spectral characteristics of DOM

DOM optical characteristics were obtained with a UV-Vis spectrophotometer (Hitachi U2900, Germany) and a spectrofluorometer (Hitachi F7000, Germany). Absorbance spectra were recorded from 190 to 800 nm with an increment of 1 nm and used to compute the specific ultraviolet absorbance at 254 nm (SUVA_254_) and absorption spectral slopes (Weishaar *et al*., 2003; Helms *et al*., 2008). SUVA_254_ is an indicator of aromaticity and chemical reactivity while the absorption slopes are used as proxies for DOM molecular weight.

Excitation emission matrices (EEMs) were generated with excitation wavelengths ranging from 220 to 450 nm and emission wavelengths ranging from 230 to 600 nm, both with 5 nm increments. EEMs were corrected with a MilliQ water sample and for inner filter effect using the absorbance-based method (Christmann *et al*., 1980; Murphy *et al*., 2013). Then, fluorescence, humification and freshness indices, as well as specific fluorescence intensity at various peaks were calculated as described by Hansen *et al*. (2016).

### Carbon stable isotope ratio

Stable isotope analysis of the respired CO_2_ provides information on carbon substrates metabolized by the microbial community (Fabian *et al*., 2017). The ratio of ^13^C/^12^C in the dissolved CO_2_ was measured directly in each microcosm by a membrane-inlet mass-spectrometer dissolved gas analyzer (HiCube pumping station, Pfeiffer Vacuum and Bay Instruments membrane, USA), controlled by Quickdata software. The sampling capillary was inserted through the PTFE-coated silicone septa prior to opening microcosms in order to avoid gas leakages. Concentrations of ^12^CO_2_ and ^13^CO_2_ were obtained from the ion currents sequentially recorded at mass to charge (m/z) ratios of 44 and 45, respectively.

For the carbon stable isotope ratio in POC and DOC, 100 ml of water was passed through GF75 filters (Advantec, USA) and both the flow-through and the filter (all five replicates were pooled onto one filter to have enough material for subsequent analyzes) were collected and freeze-dried. Moreover, we dried 1 mg of acid-killed daphnia and approximately 1 mg of extracted humic matter for carbon stable isotope ratio analysis. The samples were analyzed with an Elemental Analyzer (Thermo Flash EA 2000), coupled to a continuous-flow isotope ratio mass spectrometer (Thermo Finnigan Delta V) via an open split interface (Thermo Finnigan Conflow IV) in the IRMS Laboratory of the Leibniz Institute for Baltic Sea Research Warnemünde (Germany), and with a PDZ Europa ANCA-GSL elemental analyzer interfaced to a PDZ Europa 20-20 isotope ratio mass spectrometer (Sercon Ltd., Cheshire, UK) at the UC Davis Stable Isotope Facility (USA).

### Decomposition calculations

To test for a possible priming effect, we used an addition model as in Hannides and Aller (2016). Initial and final total organic carbon concentrations in the humic control, i.e. TOC_i_ (**H**) and TOC_f_ (H), respectively, in the carcass control, i.e. TOC_i_ (D) and TOC_f_ (D), and in the blank, i.e. TOC_i_ (B) and TOC_f_ (B), respectively, were used to predict the degradation of total organic carbon (ΔTOC) in the **HD** microcosm. Thereby, we assumed that the degradation rates of carbon sources observed in the control microcosms are conserved when humic matter and carcasses are incubated together. The predicted ΔTOC(H+D) was compared with the measured ΔTOC(HD) to detect any enhanced degradation of organic carbon caused by a potential “priming effect”. Similarly, data on CO_2_ concentration and isotope ratio (^13^CO_2_/^12^CO_2_) in the microcosms was used to calculate respiration of recalcitrant and labile carbon pools. We assumed that if a priming effect is present, there should be a difference in the observed isotope ratio between measured and predicted CO_2_, based on the sum of control values.

The carcass carbon fraction (F_c_) in DOC, POC and CO_2_ was calculated via a stable isotope mixing model (Hopkins and Ferguson, 2012). We also applied Keeling plot analyses of dissolved CO_2_ (Pataki *et al*., 2003) to estimate the δ^13^C of the carbon source respired in the respective microcosms. Descriptive calculation formulas are available in the Supplementary Methods.

### Statistical analyses

For alpha-diversity calculations of bacterial communities, individual samples were rarefied to the lowest number of reads in a sample (10080), with 10 iterations per sample using QIIME (Caporaso *et al*., 2010). Shannon biodiversity index was calculated to estimate richness and evenness of the microcosm communities. Beta-diversity of bacterial communities was calculated by using the Bray-Curtis similarity coefficients (Bray and Curtis, 1957) using QIIME.

All analyses described below were run via R version 3.3.3 (R Development Core Team, 2006). R package *vegan* (vegan: Community Ecology Package, 2017, https://CRAN.R-project.org/package=vegan) was used to carry out principal coordinate analysis (PCoA) and to calculate ANOSIM and SIMPER (Clarke, 1993).

Bacterial community composition was linked to DOC characteristics which significantly differentiated microcosms. We used the least absolute shrinkage and selection operator (LASSO) to select genera that were influenced by a variation in DOM composition (Traving *et al*., 2016). Before designing the LASSO models, we summed the abundances of OTUs belonging to the same genus and performed a centered log-ratio transformation (Gloor and Reid, 2016) of the absolute abundances of all genera. To visualize the outcome, we depicted the selected genera in two different networks using Cytoscape software (Shannon *et al*., 2003).

All differences between treatments were tested using a one-way analysis of variance (ANOVA) and post-hoc Tukey tests. Pairwise comparisons between initial and final parameters of microcosms and between two treatments were done using Student’s t-test and Wilcoxon rank sum test. Normality was checked by Shapiro-Wilks tests when necessary.

## Acknowledgements

This research was funded by German Science Foundation (GR 1540/29-1, GR1549/23-1) and the Russian Foundation for Basic Research (No. 16-54-12048), and partly supported by Russian Federal Tasks of Fundamental Research (project No. 51.1.1) and by the Council on grants from the President of the Russian Federation for support of Leading Scientific Schools (grant NSh-9249.2016.5). OVK was supported by Michail-Lomonosov-Programme-Linie A, 2015 (57180771) funded by the Ministry of Education and Science of the Russian Federation and the German Academic Exchange Service (DAAD). The authors thank Uta Mallok for DOC and phosphate concentration measurements. We are grateful to Maren Voss and Iris Liskow in the IRMS Laboratory of the Leibniz Institute for Baltic Sea Research Warnemünde (Germany) for stable isotope analysis. We greatly appreciate the useful advices of Jenny Fabian and Isabell Klawonn on stable isotope sample preparation and data handling.

The authors declare no conflict of interests.

## References

Adams, H.E., Crump, B.C., and Kling, G.W. (2010) Temperature controls on aquatic bacterial production and community dynamics in arctic lakes and streams. Environ Microbiol 12: 1319–1333.

Amado, A.M., Cotner, J.B., Cory, R.M., Edhlund, B.L., and McNeill, K. (2015) Disentangling the interactions between photochemical and bacterial degradation of dissolved organic matter: amino acids play a central role. Microb Ecol 69: 554–566.

Attermeyer, K., Hornick, T., Kayler, Z.E., Bahr, A., Zwirnmann, E., Grossart, H.-P., and Premke, K. (2014) Enhanced bacterial decomposition with increasing addition of autochthonous to allochthonous carbon without any effect on bacterial community composition. Biogeosciences 11: 1479–1489.

Attermeyer, K., Tittel, J., Allgaier, M., Frindte, K., Wurzbacher, C., Hilt, S., et al. (2015) Effects of light and autochthonous carbon additions on microbial turnover of allochthonous organic carbon and community composition. Microb Ecol 69: 361–371.

Battin, T.J., Luyssaert, S., Kaplan, L.A., Aufdenkampe, A.K., Richter, A., and Tranvik, L.J. (2009) The boundless carbon cycle. Nat Geosci 2: 598–600.

Bell, K.S., Philp, J.C., Aw, D.W.J., and Christofi, N. (1998) The genus *Rhodococcus*. J Appl Microbiol 85: 195–210.

Bengtsson, M.M., Wagner, K., Burns, N.R., Herberg, E.R., Wanek, W., Kaplan, L.A., and Battin, T.J. (2014) No evidence of aquatic priming effects in hyporheic zone microcosms. Sci Rep 4: 5187.

Bianchi, T.S. (2011) The role of terrestrially derived organic carbon in the coastal ocean: A changing paradigm and the priming effect. Proc Natl Acad Sci 108: 19473–19481.

Bianchi, T.S., Thornton, D.C.O., Yvon-Lewis, S.A., King, G.M., Eglinton, T.I., Shields, M.R., et al. (2015) Positive priming of terrestrially derived dissolved organic matter in a freshwater microcosm system. Geophys Res Lett 42: 5460–5467.

Bickel, S.L., and Tang, K.W. (2010) Microbial decomposition of proteins and lipids in copepod versus rotifer carcasses. Mar Biol 157: 1613–1624.

Blanchet, M., Pringault, O., Bouvy, M., Catala, P., Oriol, L., Caparros, J., et al. (2015) Changes in bacterial community metabolism and composition during the degradation of dissolved organic matter from the jellyfish *Aurelia aurita* in a Mediterranean coastal lagoon. Environ Sci Pollut Res 22: 13638–13653.

Blanchet, M., Pringault, O., Panagiotopoulos, C., Lefevre, D., Charriere, B., Ghiglione, J.-F., et al. (2017) When riverine dissolved organic matter (DOM) meets labile DOM in coastal waters: changes in bacterial community activity and composition. Aquat Sci 79: 27–43.

Bray, J.R., and Curtis, J.T. (1957) An ordination of the upland forest communities of Southern Wisconsin. Ecol Monogr 27: 325–349.

Caporaso, J.G., Kuczynski, J., Stombaugh, J., Bittinger, K., Bushman, F.D., Costello, E.K., et al. (2010) QIIME allows analysis of high-throughput community sequencing data. Nat Methods 7: 335–336.

Catalán, N., Kellerman, A.M., Peter, H., Carmona, F., and Tranvik, L.J. (2015) Absence of a priming effect on dissolved organic carbon degradation in lake water. Limnol Oceanogr 60: 159–168.

Christmann, D.R., Crouch, S.R., Holland, J.F., and Timnick, A. (1980) Correction of right-angle molecular fluorescence measurements for absorption of fluorescence radiation. Anal Chem 52: 291–295.

Clarke, K.R. (1993) Non-parametric multivariate analyses of changes in community structure. Austral Ecol 18: 117–143.

Dorado-García, I., Syväranta, J., Devlin, S.P., Medina-Sánchez, J.M., and Jones, R.I. (2015) Experimental assessment of a possible microbial priming effect in a humic boreal lake. Aquat Sci 1–12.

Doronina, N., Kaparullina, E., and Trotsenko, Y. (2014) The Family Methylophilaceae. In Rosenberg, E., DeLong, E.F., Lory, S., Stackebrandt, E., and Thompson, F. (eds), The Prokaryotes: Alphaproteobacteria and Betaproteobacteria. Springer Berlin Heidelberg, Berlin, Heidelberg, pp. 869–880.

Dubovskaya, O.P., Gladyshev, M.I., Gubanov, V.G., and Makhutova, O.N. (2003) Study of non-consumptive mortality of Crustacean zooplankton in a Siberian reservoir using staining for live/dead sorting and sediment traps. Hydrobiologia 504: 223–227.

Dubovskaya, O.P., Tolomeev, A.P., Kirillin, G., Buseva, Z., Tang, K.W., and Gladyshev, M.I. (2018) Effects of water column processes on the use of sediment traps to measure zooplankton non-predatory mortality: a mathematical and empirical assessment. J Plankton Res 40: 91–106.

Elliott, D.T., Harris, C.K., and Tang, K.W. (2010) Dead in the water: The fate of copepod carcasses in the York River estuary, Virginia. Limnol Oceanogr 55: 1821–1834.

Evtushenko, L.I. (2015) Marmoricola. In Bergey’s Manual of Systematics of Archaea and Bacteria. John Wiley & Sons, Ltd.

Fabian, J., Zlatanovic, S., Mutz, M., and Premke, K. (2017) Fungal–bacterial dynamics and their contribution to terrigenous carbon turnover in relation to organic matter quality. ISME J 11: 415–425.

Fonte, E.S., Amado, A.M., Meirelles-Pereira, F., Esteves, F.A., Rosado, A.S., and Farjalla, V.F. (2013) The combination of different carbon sources enhances bacterial growth efficiency in aquatic ecosystems. Microb Ecol 66: 871–878.

Gloor, G.B., and Reid, G. (2016) Compositional analysis: a valid approach to analyze microbiome high-throughput sequencing data. Can J Microbiol 62: 692–703.

Glud, R.N., Grossart, H.-P., Larsen, M., Tang, K.W., Arendt, K.E., Rysgaard, S., et al. (2015) Copepod carcasses as microbial hot spots for pelagic denitrification. Limnol Oceanogr 60: 2026–2036.

Gómez-Consarnau, L., Lindh, M.V., Gasol, J.M., and Pinhassi, J. (2012) Structuring of bacterioplankton communities by specific dissolved organic carbon compounds. Environ Microbiol 14: 2361–2378.

Gontikaki, E., Thornton, B., Huvenne, V.A.I., and Witte, U. (2013) Negative priming effect on organic matter mineralisation in NE Atlantic slope sediments. PLoS ONE 8: e67722.

Gontikaki, E., Thornton, B., Cornulier, T., and Witte, U. (2015) Occurrence of priming in the degradation of lignocellulose in marine sediments. PLoS ONE 10: e0143917.

Goto, M. (2015) Rhizobacter. In, Bergey’s Manual of Systematics of Archaea and Bacteria. John Wiley & Sons, Ltd.

Grinhut, T., Hadar, Y., and Chen, Y. (2007) Degradation and transformation of humic substances by saprotrophic fungi: processes and mechanisms. Fungal Biol Rev. 21: 179–189.

Grossart, H.-P. (2010) Ecological consequences of bacterioplankton lifestyles: changes in concepts are needed. Environ Microbiol Rep 2: 706–714.

Grossart, H.-P. and Rojas-Jimenez, K. (2016) Aquatic fungi: targeting the forgotten in microbial ecology. Curr Opin Microbiol 31: 140–145.

Grossart, H.-P., Tang, K.W., Kiørboe, T., and Ploug, H. (2007) Comparison of cell-specific activity between free-living and attached bacteria using isolates and natural assemblages. FEMS Microbiol Lett 266: 194–200.

Grossart, H.-P., Jezbera, J., Hornak, K., Hutalle, K.M.L., Buck, U., and Simek, K. (2008) Top-down and bottom-up induced shifts in bacterial abundance, production and community composition in an experimentally divided humic lake. Environ Microbiol 10: 635–652.

Guenet, B., Danger, M., Harrault, L., Allard, B., Jauset-Alcala, M., Bardoux, G., et al. (2013) Fast mineralization of land-born C in inland waters: first experimental evidences of aquatic priming effect. Hydrobiologia 721: 35–44.

Hannides, A.K. and Aller, R.C. (2016) Priming effect of benthic gastropod mucus on sedimentary organic matter remineralization. Limnol Oceanogr 61: 1640–1650.

Hansen, A.M., Kraus, T.E.C., Pellerin, B.A., Fleck, J.A., Downing, B.D., and Bergamaschi, B.A. (2016) Optical properties of dissolved organic matter (DOM): Effects of biological and photolytic degradation. Limnol Oceanogr 61: 1015–1032.

Heathcote, A., Filstrup, C., Kendall, D., and Downing, J. (2016) Biomass pyramids in lake plankton: influence of Cyanobacteria size and abundance. Inland Waters 6: 250-257.

Helms, J.R., Stubbins, A., Ritchie, J.D., Minor, E.C., Kieber, D.J., and Mopper, K. (2008) Absorption spectral slopes and slope ratios as indicators of molecular weight, source, and photobleaching of chromophoric dissolved organic matter. Limnol Oceanogr 53: 955–969.

Henson, M.W., Hanssen, J., Spooner, G., Flemming, P., Pukonen, M., Stahr, F., and Thrash, J.C. (2016) Microbial regime changes and indicators of eutrophication on the Mississippi River identified via a human-powered 2900 km transect. bioRxiv. https://doi.org/10.1101/091512

Hoikkala, L., Tammert, H., Lignell, R., Eronen-Rasimus, E., Spilling, K., and Kisand, V. (2016) Autochthonous dissolved organic matter drives bacterial community composition during a bloom of filamentous Cyanobacteria. Front Mar Sci 3: 111.

Hopkins, J.B. and Ferguson, J.M. (2012) Estimating the diets of animals using stable isotopes and a comprehensive Bayesian mixing model. PLoS ONE 7: e28478.

Hotchkiss, E.R., Hall, R.O., Baker, M.A., Rosi-Marshall, E.J., and Tank, J.L. (2014) Modeling priming effects on microbial consumption of dissolved organic carbon in rivers. J Geophys Res Biogeosciences 119: 982–995.

Huber, S.A., Balz, A., Abert, M., and Pronk, W. (2011) Characterisation of aquatic humic and non-humic matter with size-exclusion chromatography-organic carbon detection-organic nitrogen detection (LC-OCD-OND). Water Res 45: 879–885.

Hutalle-Schmelzer, K.M.L., Zwirnmann, E., Krüger, A., and Grossart, H.-P. (2010) Enrichment and cultivation of pelagic bacteria from a humic lake using phenol and humic matter additions. FEMS Microbiol Ecol 72: 58–73.

Johansen, J.E., Binnerup, S.J., Kroer, N., and Mølbak, L. (2005) *Luteibacter rhizovicinus gen. nov*., *sp. nov*., a yellow-pigmented gammaproteobacterium isolated from the rhizosphere of barley (*Hordeum vulgare* L.). Int J Syst Evol Microbiol 55: 2285–2291.

Kämpfer, P. (2015) Sphingobacteriales ord. nov. In Bergey’s Manual of Systematics of Archaea and Bacteria. John Wiley & Sons, Ltd.

Kirillin, G., Grossart, H.-P., and Tang, K.W. (2012) Modeling sinking rate of zooplankton carcasses: Effects of stratification and mixing. Limnol Oceanogr 57: 881–894.

Kisand, V., Gebhardt, S., Rullkötter, J., and Simon, M. (2013) Significant bacterial transformation of riverine humic matter detected by pyrolysis GC–MS in serial chemostat experiments. Mar Chem 149: 23–31.

Klindworth, A., Pruesse, E., Schweer, T., Peplies, J., Quast, C., Horn, M., and Glöckner, F.O. (2013) Evaluation of general 16S ribosomal RNA gene PCR primers for classical and next-generation sequencing-based diversity studies. Nucleic Acids Res 41: e1.

Kuehn, K.A., Francoeur, S.N., Findlay, R.H., and Neely, R.K. (2014) Priming in the microbial landscape: periphytic algal stimulation of litter-associated microbial decomposers. Ecology 95: 749–762.

Kuykendall, L.D. (2005) Order VI. Rhizobiales ord. nov. In, Bergey’s Manual of Systematics of Archaea and Bacteria. John Wiley & Sons, Ltd.

Landa, M., Blain, S., Christaki, U., Monchy, S., and Obernosterer, I. (2016) Shifts in bacterial community composition associated with increased carbon cycling in a mosaic of phytoplankton blooms. ISME J 10: 39–50.

Lindh, M.V., Lefébure, R., Degerman, R., Lundin, D., Andersson, A., and Pinhassi, J. (2015) Consequences of increased terrestrial dissolved organic matter and temperature on bacterioplankton community composition during a Baltic Sea mesocosm experiment. AMBIO 44: 402–412.

McBride, M.J. (2014) The Family Flavobacteriaceae. In Rosenberg, E., DeLong, E.F., Lory, S., Stackebrandt, E., and Thompson, F. (eds), The Prokaryotes: other major lineages of Bacteria and the Archaea. Springer Berlin Heidelberg, Berlin, Heidelberg, pp. 643–676.

Meziti, A., Kormas, K.A., Moustaka-Gouni, M., and Karayanni, H. (2015) Spatially uniform but temporally variable bacterioplankton in a semi-enclosed coastal area. Syst Appl Microbiol 38: 358–367.

Murphy, K.R., Stedmon, C.S., Graeber, D., and Bro, R. (2013) Fluorescence spectroscopy and multi-way techniques. PARAFAC. Anal Methods 5: 6557–6566.

Nercessian, O., Noyes, E., Kalyuzhnaya, M.G., Lidstrom, M.E., and Chistoserdova, L. (2005) Bacterial populations active in metabolism of C1 compounds in the sediment of Lake Washington, a freshwater lake. Appl Environ Microbiol 71: 6885–6899.

Newton, R.J., Jones, S.E., Eiler, A., McMahon, K.D., and Bertilsson, S. (2011) A guide to the natural history of freshwater lake bacteria. Microbiol Mol Biol Rev 75: 14–49.

van Nugteren, P., Moodley, L., Brummer, G.-J., Heip, C.H.R., Herman, P.M.J., and Middelburg, J.J. (2009) Seafloor ecosystem functioning: the importance of organic matter priming. Mar Biol 156: 2277–2287.

O’Sullivan, L.A., Rinna, J., Humphreys, G., Weightman, A.J., and Fry, J.C. (2005) *Fluviicola taffensis gen. nov., sp. nov.*, a novel freshwater bacterium of the family *Cryomorphaceae* in the phylum ‘*Bacteroidetes*’. Int J Syst Evol Microbiol 55: 2189–2194.

Pace, M.L., Cole, J.J., Carpenter, S.R., Kitchell, J.F., Hodgson, J.R., Van de Bogert, M.C., et al. (2004) Whole-lake carbon-13 additions reveal terrestrial support of aquatic food webs. Nature 427: 240–243.

Pataki, D.E., Ehleringer, J.R., Flanagan, L.B., Yakir, D., Bowling, D.R., Still, C.J., et al. (2003) The application and interpretation of Keeling plots in terrestrial carbon cycle research. Glob Biogeochem Cycles 17: 1022.

Rocker, D., Kisand, V., Scholz-Böttcher, B., Kneib, T., Lemke, A., Rullkötter, J., and Simon, M. (2012a) Differential decomposition of humic acids by marine and estuarine bacterial communities at varying salinities. Biogeochemistry 111: 331–346.

Rocker, D., Brinkhoff, T., Grüner, N., Dogs, M., and Simon, M. (2012b) Composition of humic acid-degrading estuarine and marine bacterial communities. FEMS Microbiol Ecol 80: 45–63.

Rojas-Jimenez, K., Fonvielle, J.A., Ma, H., and Grossart, H.-P. (2017) Transformation of humic substances by the freshwater Ascomycete Cladosporium sp. Limnol Oceanogr 62: 1955-1962.

Schloss, P.D., Westcott, S.L., Ryabin, T., Hall, J.R., Hartmann, M., Hollister, E.B., et al. (2009) Introducing mothur: open-source, platform-independent, community-supported software for describing and comparing microbial communities. Appl Environ Microbiol 75: 7537–7541.

Schmidt, M.W.I., Torn, M.S., Abiven, S., Dittmar, T., Guggenberger, G., Janssens, I.A., et al. (2011) Persistence of soil organic matter as an ecosystem property. Nature 478: 49–56.

Segers, P., Vancanneyt, M., Pot, B., Torck, U., Hoste, B., Dewettinck, D., et al. (1994) Classification of *Pseudomonas diminuta* Leifson and Hugh 1954 and *Pseudomonas vesicularis* Busing, Doll, and Freytag 1953 in *Brevundimonas gen. nov*. as *Brevundimonas diminuta comb. nov*. and *Brevundimonas vesicularis comb. nov*., respectively.

Shannon, P., Markiel, A., Ozier, O., Baliga, N.S., Wang, J.T., Ramage, D., et al. (2003) Cytoscape: a software environment for integrated models of biomolecular interaction networks. Genome Res 13: 2498–2504.

Shoemaker, K.M. and Moisander, P.H. (2015) Microbial diversity associated with copepods in the North Atlantic subtropical gyre. FEMS Microbiol Ecol 91: fiv064-fiv064.

Song, J., Choo, Y.-J., and Cho, J.-C. (2008) *Perlucidibaca piscinae gen. nov., sp. nov*., a freshwater bacterium belonging to the family *Moraxellaceae*. Int J Syst Evol Microbiol 58: 97–102.

Spencer, R.G.M., Aiken, G.R., Butler, K.D., Dornblaser, M.M., Striegl, R.G., and Hernes, P.J. (2009) Utilizing chromophoric dissolved organic matter measurements to derive export and reactivity of dissolved organic carbon exported to the Arctic Ocean: A case study of the Yukon River, Alaska. Geophys Res Lett 36: L06401, doi:10.1029/2008GL036831.

Steen, A.D., Quigley, L.N.M., and Buchan, A. (2015) Evidence for the priming effect in a planktonic estuarine microbial community. Front Mar Sci 3: 6.

Steinberg, D.K. and Landry, M.R. (2017) Zooplankton and the ocean carbon cycle. Annu Rev Mar Sci 9: 413–444.

Stubbins, A., Spencer, R.G.M., Chen, H., Hatcher, P.G., Mopper, K., Hernes, P.J., et al. (2010) Illuminated darkness: Molecular signatures of Congo River dissolved organic matter and its photochemical alteration as revealed by ultrahigh precision mass spectrometry. Limnol Oceanogr 55: 1467–1477.

Tang, K.W., Hutalle, K.M.L., and Grossart, H. (2006) Microbial abundance, composition and enzymatic activity during decomposition of copepod carcasses. Aquat Microb Ecol 45: 219–227.

Tang, K., Dziallas, C., Hutalle-Schmelzer, K., and Grossart, H.-P. (2009) Effects of food on bacterial community composition associated with the copepod *Acartia tonsa* Dana. Biol Lett 5: 549.

Tang, K.W., Turk, V., and Grossart, H. (2010) Linkage between crustacean zooplankton and aquatic bacteria. Aquat Microb Ecol 61: 261–277.

Tang, K.W., Gladyshev, M.I., Dubovskaya, O.P., Kirillin, G., and Grossart, H.-P. (2014) Zooplankton carcasses and non-predatory mortality in freshwater and inland sea environments. J Plankton Res 36: 597–612.

Tranvik, L.J. and Kokalj, S. (1998) Decreased biodegradability of algal DOC due to interactive effects of UV radiation and humic matter. Aquat Microb Ecol 14: 301–307.

Tranvik, L.J., Downing, J.A., Cotner, J.B., Loiselle, S.A., Striegl, R.G., Ballatore, T.J., et al. (2009) Lakes and reservoirs as regulators of carbon cycling and climate. Limnol Oceanogr 54: 2298–2314.

Traving, S.J., Bentzon-Tilia, M., Knudsen-Leerbeck, H., Mantikci, M., Hansen, J.L.S., Stedmon, C.A., et al. (2016) Coupling bacterioplankton populations and environment to community function in coastal temperate waters. Front Microbiol 7: 1533.

Trusova, M.Y., Kolmakova, O.V., and Gladyshev, M.I. (2012) Seasonal features of consumption of lysine by uncultivated bacterial plankton of eutrophic water reservoir. Contemp Probl Ecol 5: 391–398.

Vachon, D., Prairie, Y.T., Guillemette, F., and del Giorgio, P.A. (2017) Modeling allochthonous dissolved organic carbon mineralization under variable hydrologic regimes in boreal lakes. Ecosystems 20: 781–795.

Ward, N.D., Keil, R.G., Medeiros, P.M., Brito, D.C., Cunha, A.C., Dittmar, T., et al. (2013) Degradation of terrestrially derived macromolecules in the Amazon River. Nat Geosci 6: 530–533.

Weishaar, J.L., Aiken, G.R., Bergamaschi, B.A., Fram, M.S., Fujii, R., and Mopper, K. (2003) Evaluation of specific ultraviolet absorbance as an indicator of the chemical composition and reactivity of dissolved organic carbon. Environ Sci Technol 37: 4702–4708.

